# A Cell Type-Specific Role for Tubb6 in Ciliogenesis of *Xenopus* Epidermal Multiciliated Cells

**DOI:** 10.1101/2025.05.31.657203

**Authors:** Xiaolu Xu, Jean Ross, Fiona Clark, Shuo Wei, Jian Sun

## Abstract

Cilia are microtubule-based organelles found on the surface of most eukaryotic cells. These microtubules are composed of α- and β-tubulin heterodimers, and different tubulin isotypes can confer distinct properties to microtubules. Despite their importance, the contribution of individual tubulin isotype to cilia formation and function remains largely unexplored in vertebrates. Here, we identify a critical role for the β-tubulin isotype Tubb6 in the formation of motile cilia in *Xenopus* epidermal multiciliated cells (MCCs). *Tubb6* mRNA is selectively expressed in MCCs, and its protein product localizes to ciliary axonemes. Loss of Tubb6 leads to a marked reduction in cilia number and length, resulting in defective MCC function. In contrast, mono-motile cilia in the gastrocoel roof plate are unaffected by Tubb6 depletion, suggesting a selective requirement for ciliogenesis in MCCs. Together, our findings uncover a cell type-specific role for Tubb6 in motile cilia formation and highlight the functional specialization of tubulin isotypes in vertebrate cilia assembly.

## Introduction

Cilia are microtubule-based cellular projections found on the surface of nearly all eukaryotic cells, where they serve essential roles in motility and signaling^1,2^. Based on their motility, cilia are broadly classified into non-motile and motile types. Non-motile cilia, also referred to as primary or sensory cilia, typically occur as solitary structures and act as signaling hubs that transduce extracellular cues into intracellular responses^3–5^. In contrast, motile cilia actively beat to generate extracellular directional fluid flow and are found either in large numbers on specialized epithelial cells known as MCCs or as solitary structures referred to as mono-motile cilia^6–8^. MCC-derived motile cilia are found in the brain ependyma, respiratory tract, and reproductive ducts, where they mediate fluid movement across epithelial surfaces^6^. Mono-motile cilia are present in the embryonic node of mammals, the gastrocoel roof plate (GRP) of *Xenopus*, and Kupffer’s vesicle in teleost fishes^9^, where they are critical for establishing left– right asymmetry during development^10^. Dysfunction of motile cilia can lead to a range of disorders known as motile ciliopathies, including chronic respiratory infections, hydrocephalus and situs inversus^6,11,12^. Structurally, the core of a cilium—the axoneme—is a microtubule-based scaffold. Axonemes in MCCs typically exhibit a “9+2” arrangement of microtubules, with nine outer microtubule doublets surrounding a central pair of singlets, whereas mono-motile cilia adopt a “9+0” structure lacking the central pair^1,13^.

Microtubules are cylindrical polymers composed of α-/β-tubulin heterodimers arranged into protofilaments. These cytoskeletal filaments are essential for numerous cellular processes, including cell division, vesicle trafficking, and ciliogenesis^14,15^. The functional diversity of microtubules is modulated by the expression of distinct tubulin isotypes and their post-translational modifications, collectively referred to as the “tubulin code.” While some tubulin isotypes exhibit functional redundancy, others play specialized roles in specific tissues and organelles^16,17^. Unlike cytoplasmic microtubules, which are highly dynamic and prone to rapid depolymerization, microtubules within cilia are relatively stable, reflecting their specialized structural roles^18,19^. Evidence from invertebrate models supports the importance of tubulin isotypes in ciliary architecture. In *C. elegans*, the α-tubulin isotype TBA-6 is expressed specifically in cephalic male neurons, where it’s indispensable for constructing the ciliary microtubule doublets^20^. In *Drosophila*, the testis specific β-tubulin isotype β2 is essential for axoneme assembly in sperm flagella and cannot be replaced by other isotypes^21^. In vertebrates, however, the functional roles of tubulin isotypes in cilia remain largely unexplored. Recent studies identified the β-tubulin isotype Tubb4b as essential for axoneme assembly in both humans and mice, with pathogenic mutations linked to ciliopathies^22–24^.

Tubb6 is a class V β-tubulin isotype previously implicated in microtubule organization during muscle differentiation and regeneration^25^. It also plays key roles in modulating microtubule dynamics and actin networks in osteoclasts, contributing to bone homeostasis^26^. Mutations in *Tubb6* have been associated with congenital cranial dysinnervation disorders (CCDDs), a developmental disorder characterized by congenital facial palsy^27,28^. However, whether Tubb6 contributes to ciliogenesis has not been investigated. Intriguingly, *Tubb6* expression is upregulated in *Tubb4b*-deficient mice^23^, suggesting potential compensatory or redundant functions in ciliary assembly.

In this study, we examined the role of Tubb6 in vertebrate motile cilia using *Xenopus* embryos as a model. We found that Tubb6 is specifically expressed in MCCs of the *Xenopus* mucociliary epidermis, and its protein product localizes to the axoneme of motile cilia. Disruption of Tubb6 function resulted in a significant decrease in both cilia number and length, leading to impaired MCC activity. Notably, mono-motile cilia in the GRP remained unaffected, indicating a selective requirement for Tubb6 in MCC ciliogenesis. These findings reveal a cell type-specific role for Tubb6 in motile cilia assembly and emphasize the functional specialization of tubulin isotypes in vertebrate ciliogenesis.

## Results

### Tubb6 is required for motile ciliary function in *Xenopus*

In *Xenopus* embryos, the mucociliary epidermis contains hundreds of MCCs, whose coordinated beating generates directional fluid flow that allows the embryos to glide on a surface – an indicator of proper motile ciliary function (Fig. 1A). To investigate the role of Tubb6, a translation-blocking morpholino (MO) targeting *Tubb6* (MO-1) was injected into one-cell stage embryos, and gliding behavior was assessed at stage 27. Embryos lacking Tubb6 exhibited a severe defect in gliding motility compared to those injected with control MO. Importantly, coinjection with MO-resistant *tubb6* mRNA fully rescued this defect, indicating that Tubb6 is essential for motile ciliary function (Fig. 1B).

**Figure 1.**
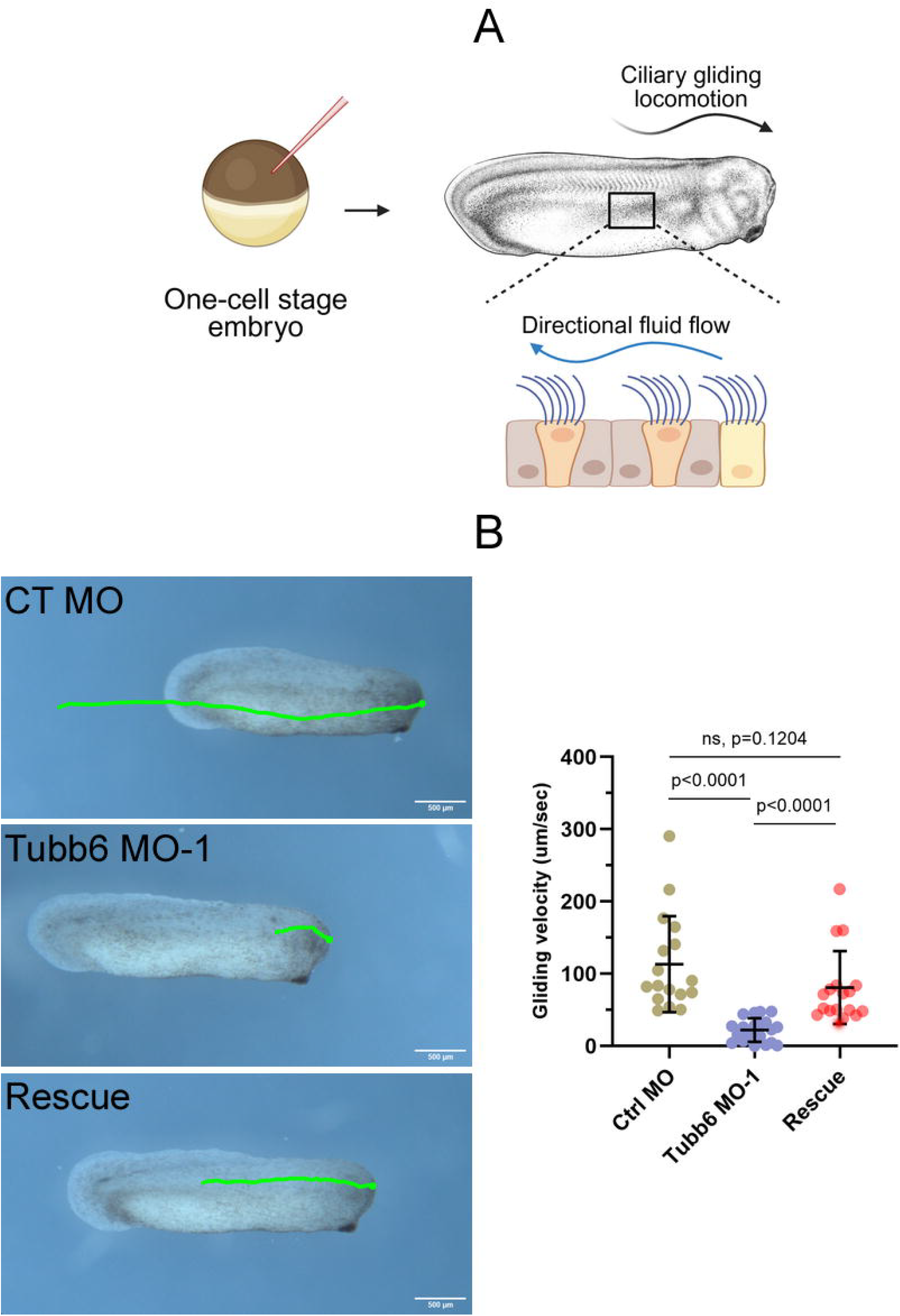
Tubb6 is required for the gilding locomotion driven by motile cilia on *Xenopus* epidermis. (A) Illustrative workflow of microinjection and gliding assays. (B) Gliding assays of St. 28 embryos. Embryos were tracked for 20Lsec by video recording, and the distance traveled by a representative embryo is indicated by green line traces, with quantification of the velocity displayed on the right. Quantitative data were collected from three biological replicates and shown as mean +/- s.d. (unpaired two-tailed t-test).

### Tubb6 is specifically expressed in MCCs and localizes to the ciliary axoneme

To determine the expression pattern of *Tubb6*, we performed whole-mount *in situ* hybridization (WISH) in *Xenopus* embryos. *Tubb6* transcripts appeared as regularly spaced puncta across the epidermis, consistent with the characteristic distribution of MCCs. Immunostaining for acetylated α-tubulin (ac-tub), a marker of ciliary axonemes, combined with WISH, further confirmed that *Tubb6* is specifically expressed in MCCs of the *Xenopus* embryonic epidermis (Fig. 2A).

**Figure 2.**
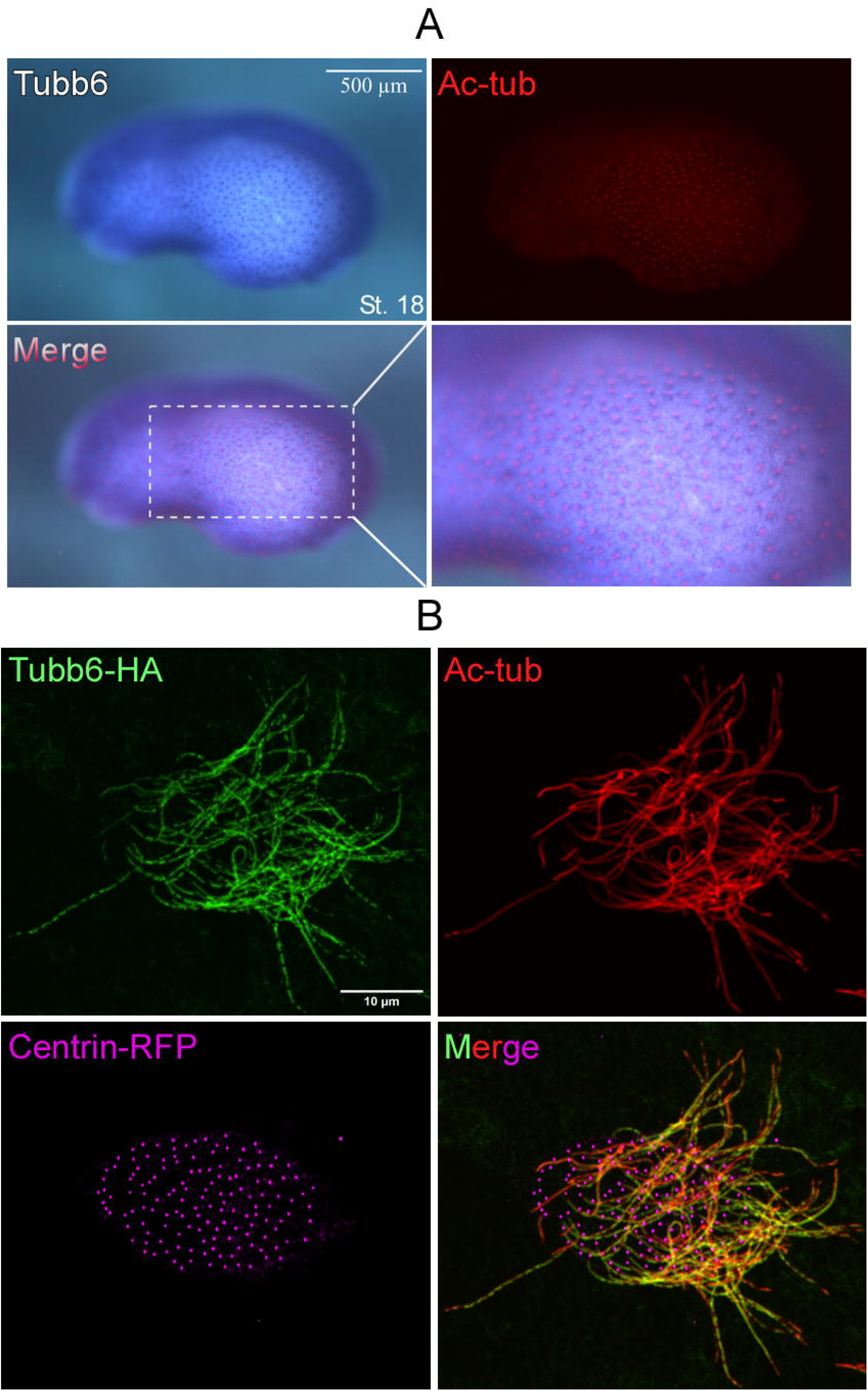
Tubb6 is localized to the ciliary axoneme in MCCs of the *Xenopus* mucociliary epidermis. (A) Embryos were fixed at St.22 and processed for WISH for *tubb6*, and then stained for ac-tub. (B) mRNA encoding HA-tagged Tubb6 was co-injected with RFP-tagged Centrin4 to label the basal body. Embryos were fixed at St. 28 and stained for HA and ac-tub.

We next examined the subcellular localization of Tubb6 protein by injecting embryos with HA-tagged *Tubb6* mRNA. Immunostaining revealed substantial co-localization of HA-Tubb6 with ac-tub along the length of the ciliary axoneme but not with RFP-tagged Centrin4, a basal body marker (Fig. 2B). This specific localization suggests that Tubb6 is a structural component of motile ciliary axonemes.

### Loss of Tubb6 disrupts ciliogenesis in MCCs

The axonemal localization of Tubb6 protein in MCCs suggested a potential role in cilia assembly. To test this, we disrupted *Tubb6* using CRISPR/Cas9 by co-injecting Cas9 mRNA and a *Tubb6*-targeting single-guide RNA (sgRNA). Genotyping confirmed efficient mutagenesis (Fig. S1). Compared to Cas9-only controls, *Tubb6* mutants exhibited significantly reduced ac-tub signal in epidermal MCCs, which was fully rescued by co-injection of wild-type *tubb6* mRNA (Fig. 3A).

**Figure 3.**
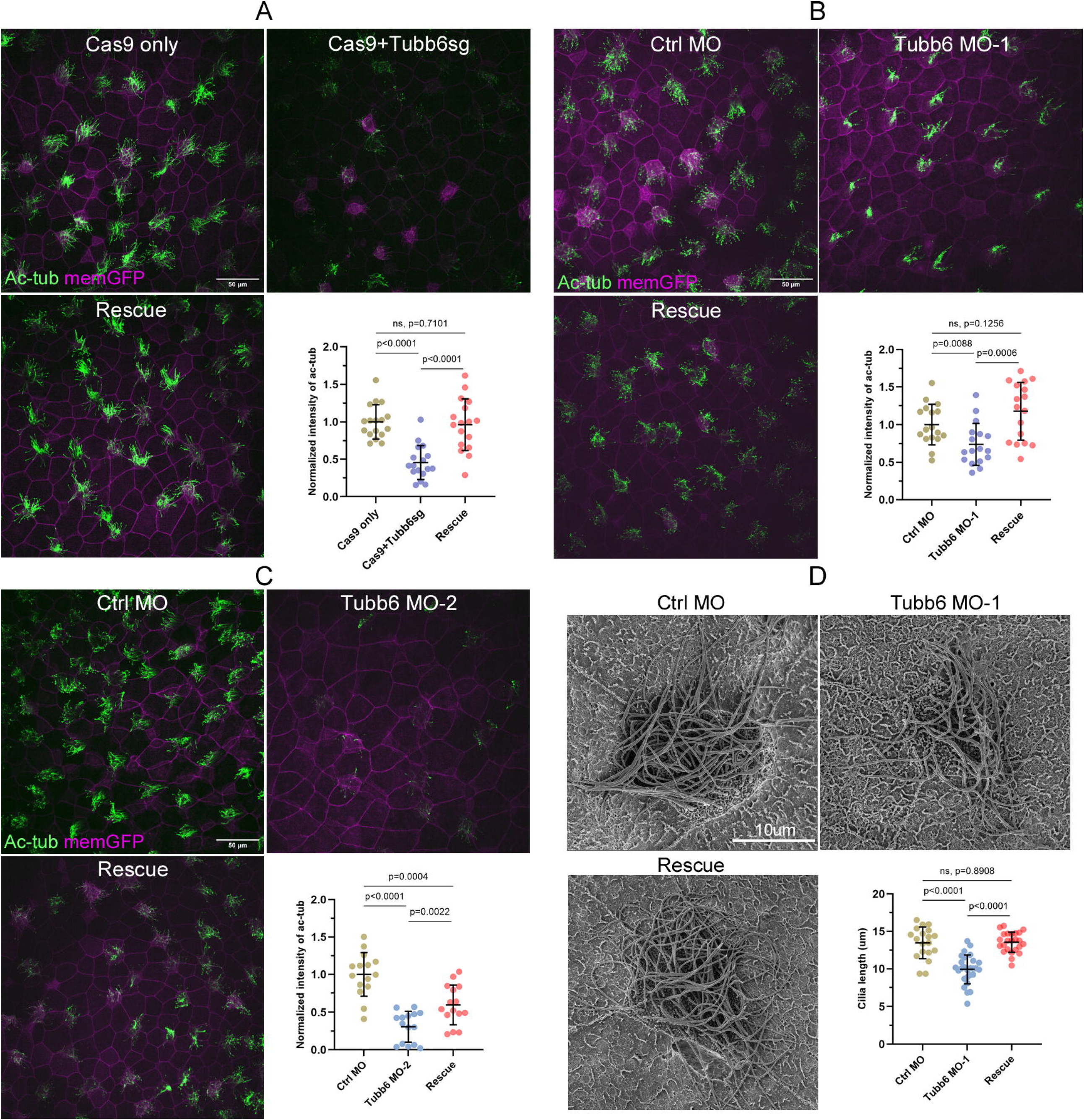
Disrupted ciliogenesis in Tubb6 crispants and morphants. (A-C) Embryos were injected with the indicated Cas9, sgRNA and MO, fixed at St. 28 and stained for ac-tub. mRNA encoding a membrane-bound form of GFP (memGFP) was co-injected as a tracer. Representative confocal images of the epidermis for each condition are shown, with quantification of ac-tub intensity on the right bottom corner. (D) SEM images of *Xenopus* embryos display impaired cilia formation in Tubb6-depleted MCCs. Quantitative data were collected from three biological replicates and shown as mean +/- s.d. (unpaired two-tailed t-test).

Consistent results were obtained using two independent translation-blocking MOs targeting distinct *Tubb6* loci. Both morphants showed diminished ac-tub staining, which could be fully (MO-1) or partially (MO-2) rescued by co-injecting MO-resistant *tubb6* mRNA (Fig. 3B and 3C), further confirming the requirement of Tubb6 for proper ciliogenesis. Scanning electron microscopy (SEM) analyses corroborated these findings, revealing a significant reduction in both cilia number and length upon Tubb6 knockdown (Fig. 3D). These results demonstrate that Tubb6 is essential for axoneme assembly and ciliary elongation in MCCs.

### Tubb6 is dispensable for mono-motile cilia formation in *Xenopus* GRP

During neurulation, GRP in the dorsal midline of *Xenopus* embryos contains ciliated cells that bear mono-motile cilia. Unlike MCCs, which possess hundreds of motile cilia, each GRP cell harbors a single cilium that generates leftward fluid flow essential for establishing left–right asymmetry. To assess *Tubb6* expression in this context, we dissected the GRP-containing dorsal tissue from stage 18 embryos and performed WISH. *Tubb6* transcripts were detected in the GRP region (Fig. 4A). To determine whether Tubb6 is required for the formation of GRP monocilia, we injected Tubb6 MO-1 into one blastomere at two-cell stage. Tubb6 knockdown did not significantly affect mono-cilia length in the GRP compared to control MO-injected embryos (Fig. 4B), indicating that Tubb6 is dispensable for GRP monocilia ciliogenesis. Together, our results demonstrate a specific role of Tubb6 in *Xenopus* epidermis MCCs, suggesting a cell type–specific requirement for this tubulin isotype during ciliogenesis.

**Figure 4.**
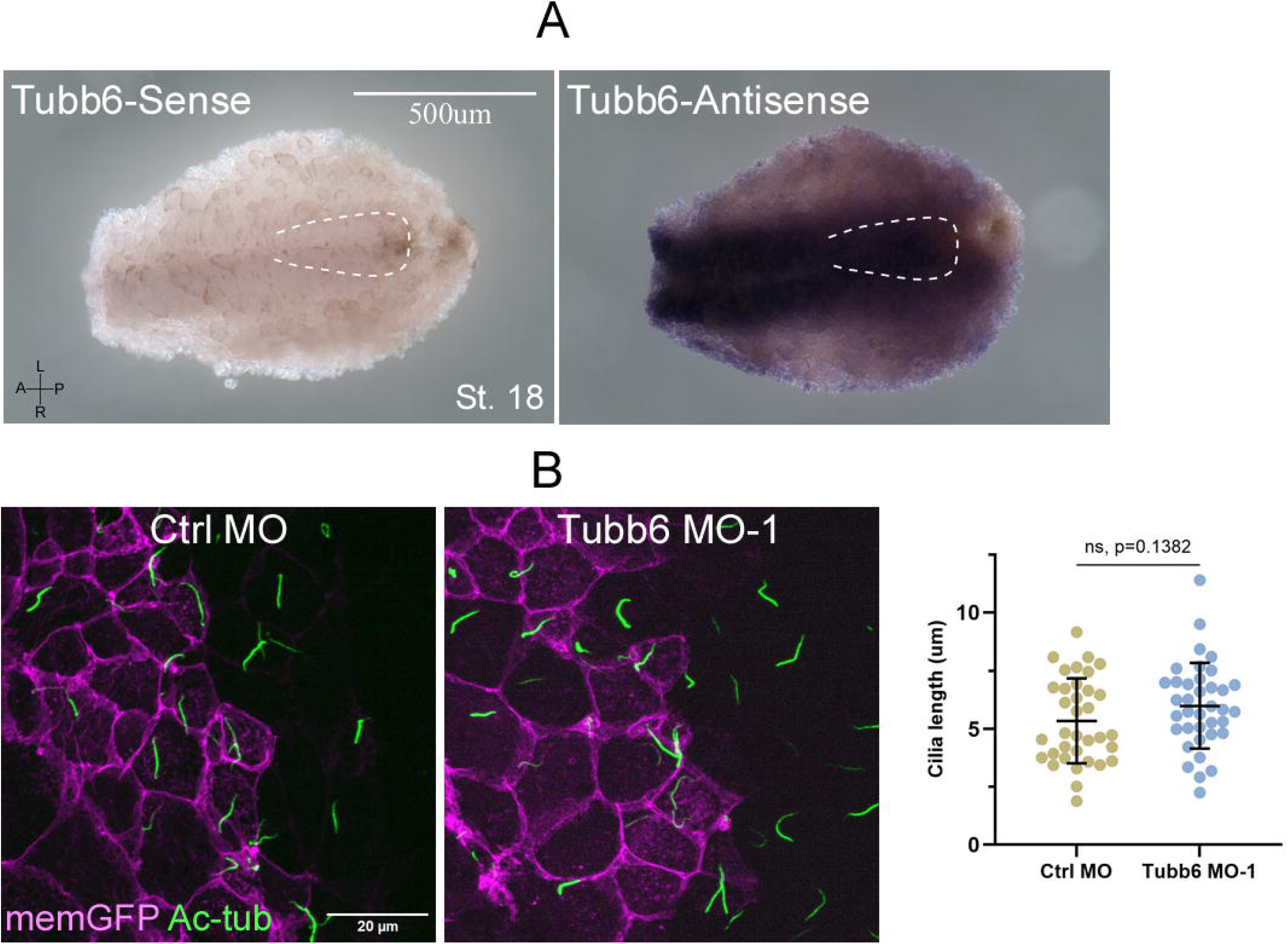
Loss of Tubb6 does not affect ciliogenesis of GRP monocilia. (A) Embryos were fixed at St. 18 and GRP-containing dorsal tissue was dissected, followed by WISH using Tubb6 sense/antisense probe. GRP region is highlighted with white dotted line. (B) Embryos were injected with indicated MO in one blastomere at two-cell stage. GRP explants were dissected at St. 18, fixed, and then stained for ac-tub. Representative confocal images of the GRP for each condition are shown, with quantification of cilia length on the right. Quantitative data were collected from two biological replicates and shown in mean +/- s.d. (unpaired two-tailed t-test).

## Discussion

Directional fluid flow generated by motile cilia is critical for various biological processes, including mucus clearance, cerebrospinal fluid circulation, and the establishment of left–right body asymmetry. Dysfunction of motile cilia results in a range of human disorders collectively known as motile ciliopathies, including primary ciliary dyskinesia, hydrocephalus, and chronic respiratory infections^11–13^. Motile cilia are specialized organelles whose core axoneme is built from α/β-tubulin heterodimers. The specific composition of tubulin isotypes influences microtubule dynamics and stability, but the extent to which individual isotypes contribute to ciliary assembly and function remains largely unclear.

In this study, we demonstrate that *tubb6* is specifically expressed in MCCs of *Xenopus* mucociliary epidermis, and that TUBB6 protein localizes exclusively to the ciliary axoneme. Notably, TUBB6 is absent from the basal body, supporting a direct role in axoneme assembly rather than in basal body anchoring or docking. Unlike TUBB4B, which has been reported to localize unevenly along the axoneme, TUBB6 is distributed along the axoneme from proximal to distal end, resembling the pattern of acetylated tubulin. This suggests that TUBB6 is incorporated throughout the axoneme and may contribute to axonemal stability or elongation. Interestingly, while some cilia exhibited continuous TUBB6 labeling, others displayed a discontinuous pattern. The underlying mechanism of this heterogeneity is unclear but could reflect the presence of tubulin heterodimers formed from different isotypes, possibly resulting in a mosaic organization within individual axonemes.

Loss of Tubb6 resulted in a pronounced defect in the gliding behavior of *Xenopus* embryos, reflecting impaired motile cilia function in MCCs. This phenotype was accompanied by a reduction in both cilia number and length, indicating that Tubb6 is essential for axoneme assembly and proper ciliogenesis. Interestingly, this requirement appears to be cell type– specific. Similar to Tubb4b, another β-tubulin isotype previously shown to be critical for MCC-derived motile cilia, Tubb6 depletion selectively disrupted MCC cilia without affecting the formation of GRP monocilia, despite its expression in the GRP. These findings highlight a shared functional specialization of Tubb6 and Tubb4b, suggesting that distinct tubulin isotypes may be differentially utilized depending on cilia type and cellular context. This cell type-specific requirement also indicates that the contribution of tubulin isotypes to ciliogenesis is context-dependent, possibly influenced by differences in ciliary architecture, protein turnover, or redundancy with other isotypes. One possible explanation is that GRP monocilia can compensate for the loss of Tubb6 through upregulation or preferential usage of other β-tubulin isotypes, maintaining sufficient tubulin availability for axoneme assembly. This is consistent with previous findings that depletion of one tubulin isotype can lead to compensatory expression of others^29,30,23^, reflecting a tightly regulated homeostasis of the cellular tubulin pool.

In summary, our findings reveal a previously uncharacterized role for Tubb6 in the assembly and function of motile cilia in MCCs. The selective requirement for Tubb6 in MCCs underscores the importance of tubulin isotype specialization in vertebrate cilia biology. These results contribute to a broader understanding of how structural diversity among tubulin subunits supports organelle-specific microtubule functions and may offer insights into tissue-specific susceptibility to tubulin-related ciliopathies.

## Methods Plasmids

Full-length cDNA clone for *Xenopus tropicalis* tubb6 (Accession: BC080942) was purchased from GE-Dharmacon. The coding sequence was then subcloned into a pCS2+ expression vector with an in-frame C-ternimal HA tag.

### Animals and embryo manipulation

Wild-type *Xenopus tropicalis* frogs were purchased from National Xenopus Resource (NXR). Methods involving live animals were carried out in accordance with the guidelines and regulations approved and enforced by the Institutional Animal Care and Use Committees at University of Delaware. Embryo were collected and injected with PLI-100A microinjectors (Harvard Apparatus). Tubb6 MO-1 (5′-CGTTAGCCCTTGCAGTGAATGGATC-3′) and MO-2 (5′-CTTGGATATGCACAATTTCCCTCAT-3′) were designed and generated by Gene Tools. The sgRNA for CRISPR/Cas9 mediated gene editing was generated as described^31^. Primer (5′-GCAGCTAATACGACTCACTATAGGTAGGAGTAGTGAGTTTCAGTTTTAGAGCTAGAAATA-3′) was used to synthesize the template for sgRNA Tubb6-sg. *In vitro* transcription was carried out to generate the sgRNA using the MEGAshortscript T7 Transcription Kit (ThermoFisher Scientific). Capped *tubb6* mRNA was transcribed using the mMessage mMachine SP6 kit (ThermoFisher Scientific). Embryos were injected with mRNAs or MO at one-cell or two-cell stage. The following RNAs were used: memGFP mRNA (100 pg), Tubb6-HA mRNA (300 pg), and Tubb6-sg (200pg). 10 ng Con MO or Tubb6 MO was injected per embryo at one-cell stage or 5 ng per blastomere at two-cell stage. Gliding assay was performed using stage 27 embryos placed on 1% agar-coated dishes. Embryo gliding was captured using a Zeiss Axio Zoom V16 fluorescence microscope with an AxioCam camera, and the movement trajectories were quantified using Manual tracking plugin of ImageJ.

### Whole-mount *in situ* hybridization

*In vitro* transcription was conducted to generate digoxigenin-labeled antisense and sense RNA probes using T3 polymerase and Sp6 polymerase, respectively. *In situ* hybridization were performed as described previously^32^. Images were taken with a Zeiss Axio Zoom V16 fluorescence microscope and an AxioCam camera.

### Immunofluorescence microscopy

Embryos were fixed with 4% formaldehyde at 4°C overnight and then permeabilized with 0.5% Triton in PBS for 10 min at room temperature. The following antibodies were used: mouse anti-acetylated tubulin-conjugated with Alexa Fluor 594 (1:500, Proteintech), Rat anti-HA (1:500, Sigma) and donkey anti-rat conjugated with Alexa Fluor 488 (1:1000, Invitrogen). The samples were mounted in ProLong Gold and imaged using Andor Dragonfly spinning disk confocal microscope.

### Electron Microscopy

The embryos were fixed with 2% paraformaldehyde, 2% glutaraldehyde in 0.1M sodium cacodylate buffer, rinsed with the buffer, and then fixed with 1% osmium tetroxide in 0.1M sodium cacodylate buffer. They were dehydrated in ascending concentrations of ethanol (25% to 100%) and then treated with hexamethyldisilazane and air dried. The dried samples were mounted on SEM stubs and sputter coated with platinum in a Leica EM ACE600 coater (Vienna, Austria). SEM images were acquired using a ThermoFisher Scientific Apreo Volumescope SEM (Hillsboro, OR).

## Statistical Analysis

All statistical significance was assessed using unpaired two-tailed t-tests, and p-values smaller than 0.05 were considered statistically significant.

## Acknowledgments

This work is supported by NIH R01 DE029802. Microscopy equipment was acquired with a shared instrumentation grant (S10 OD025165) and access was supported by NIH—NIGMS (P20 GM103446), NIGMS (P20 GM139760) and the State of Delaware.

## Reference

1. Satir, P. & Christensen, S. T. Overview of Structure and Function of Mammalian Cilia. Annual Review of Physiology 69, 377–400 (2007).

2. Nigg, E. A. & Raff, J. W. Centrioles, Centrosomes, and Cilia in Health and Disease. Cell 139, 663–678 (2009).

3. Singla, V. & Reiter, J. F. The Primary Cilium as the Cell’s Antenna: Signaling at a Sensory Organelle. Science 313, 629–633 (2006).

4. Satir, P., Pedersen, L. B. & Christensen, S. T. The primary cilium at a glance. Journal of Cell Science 123, 499–503 (2010).

5. Goetz, S. C. & Anderson, K. V. The primary cilium: a signalling centre during vertebrate development. Nat Rev Genet 11, 331–344 (2010).

6. Brooks, E. R. & Wallingford, J. B. Multiciliated Cells. Current Biology 24, R973–R982 (2014).

7. Mitchell, B., Jacobs, R., Li, J., Chien, S. & Kintner, C. A positive feedback mechanism governs the polarity and motion of motile cilia. Nature 447, 97–101 (2007).

8. Nonaka, S., Shiratori, H., Saijoh, Y. & Hamada, H. Determination of left–right patterning of the mouse embryo by artificial nodal flow. Nature 418, 96–99 (2002).

9. Essner, J. J. et al. Conserved function for embryonic nodal cilia. Nature 418, 37–38 (2002).

10. Hirokawa, N., Tanaka, Y., Okada, Y. & Takeda, S. Nodal Flow and the Generation of Left-Right Asymmetry. Cell 125, 33–45 (2006).

11. Fliegauf, M., Benzing, T. & Omran, H. When cilia go bad: cilia defects and ciliopathies. Nat Rev Mol Cell Biol 8, 880–893 (2007).

12. Wallmeier, J. et al. Motile ciliopathies. Nat Rev Dis Primers 6, 1–29 (2020).

13. Ishikawa, H. & Marshall, W. F. Ciliogenesis: building the cell’s antenna. Nat Rev Mol Cell Biol 12, 222–234 (2011).

14. Welte, M. A. Bidirectional Transport along Microtubules. Current Biology 14, R525–R537 (2004).

15. Prosser, S. L. & Pelletier, L. Mitotic spindle assembly in animal cells: a fine balancing act. Nat Rev Mol Cell Biol 18, 187–201 (2017).

16. Westermann, S. & Weber, K. Post-translational modifications regulate microtubule function. Nat Rev Mol Cell Biol 4, 938–948 (2003).

17. Janke, C. & Magiera, M. M. The tubulin code and its role in controlling microtubule properties and functions. Nat Rev Mol Cell Biol 21, 307–326 (2020).

18. Mitchison, T. & Kirschner, M. Dynamic instability of microtubule growth. Nature 312, 237–242 (1984).

19. Wloga, D., Joachimiak, E., Louka, P. & Gaertig, J. Posttranslational Modifications of Tubulin and Cilia. Cold Spring Harb Perspect Biol 9, a028159 (2017).

20. Silva, M. et al. Cell-Specific α-Tubulin Isotype Regulates Ciliary Microtubule Ultrastructure, Intraflagellar Transport, and Extracellular Vesicle Biology. Current Biology 27, 968–980 (2017).

21. Hoyle, H. D. & Raff, E. C. Two Drosophila beta tubulin isoforms are not functionally equivalent. Journal of Cell Biology 111, 1009–1026 (1990).

22. Dodd, D. O. et al. Ciliopathy patient variants reveal organelle-specific functions for TUBB4B in axonemal microtubules. Science 384, eadf5489 (2024).

23. Sanzhaeva, U. et al. TUBB4B is essential for the cytoskeletal architecture of cochlear supporting cells and motile cilia development. Commun Biol 7, 1–18 (2024).

24. Sewell, M. T., Legué, E. & Liem, K. F., Jr. Tubb4b is required for multi-ciliogenesis in the mouse. Development 151, dev201819 (2024).

25. Randazzo, D. et al. Persistent upregulation of the β-tubulin tubb6, linked to muscle regeneration, is a source of microtubule disorganization in dystrophic muscle. Hum Mol Genet 28, 1117–1135 (2019).

26. Maurin, J. et al. The Beta-Tubulin Isotype TUBB6 Controls Microtubule and Actin Dynamics in Osteoclasts. Front Cell Dev Biol 9, 778887 (2021).

27. Fazeli, W. et al. A TUBB6 mutation is associated with autosomal dominant non-progressive congenital facial palsy, bilateral ptosis and velopharyngeal dysfunction. Hum Mol Genet 26, 4055–4066 (2017).

28. Jurgens, J. A. et al. Expanding the genetics and phenotypes of ocular congenital cranial dysinnervation disorders. Genetics in Medicine 27, 101216 (2025).

29. Latremoliere, A. et al. Neuronal-Specific TUBB3 Is Not Required for Normal Neuronal Function but Is Essential for Timely Axon Regeneration. Cell Reports 24, 1865-1879.e9 (2018).

30. Fertuzinhos, S., Legué, E., Li, D. & Liem, K. F. A dominant tubulin mutation causes cerebellar neurodegeneration in a genetic model of tubulinopathy. Science Advances 8, eabf7262 (2022).

31. Nakayama, T. et al. Cas9-based genome editing in Xenopus tropicalis. Methods Enzymol 546, 355–375 (2014).

32. Wei, S. et al. ADAM13 Induces Cranial Neural Crest by Cleaving Class B Ephrins and Regulating Wnt Signaling. Developmental Cell 19, 345–352 (2010).

