## Supplemental Figure 1 for "A Cell Type-Specific Role for Tubb6 in Ciliogenesis of *Xenopus* Epidermal Multiciliated Cells"

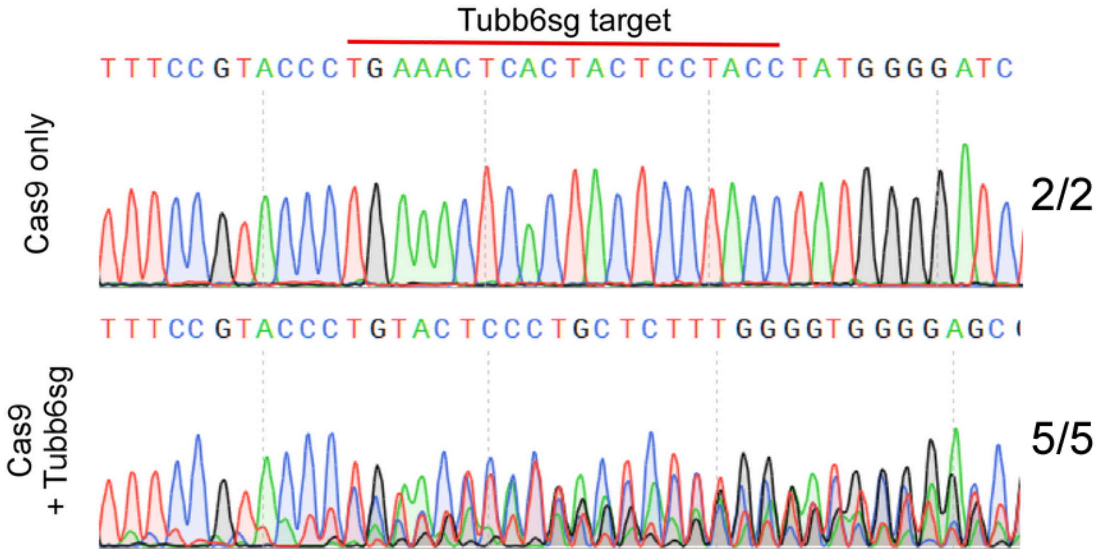


**Figure S1. Genotyping of control embryos and Tubb6 crispants.**
